# Application of chimeric antigens to paper-based diagnostics for detection of West Nile virus infections of *Crocodylus porosus –* a novel animal test case

**DOI:** 10.1101/2024.03.24.586480

**Authors:** Ryan A. Johnston, Gervais Habarugira, Jessica J. Harrison, Sally R. Isberg, Jasmin Moran, Mahali Morgan, Steven S. Davis, Lorna Melville, Christopher B. Howard, Charles S. Henry, Joanne Macdonald, Helle Bielefeldt-Ohmann, Roy A. Hall, Jody Hobson-Peters

**Author notes:** Author for Correspondence: Dr Jody Hobson-Peters School of Chemistry and Molecular Biosciences, The University of Queensland, St. Lucia, Queensland, 4072 Australia E.

## Abstract

Laboratory-based diagnostics like plaque reduction neutralization tests (PRNT) and ELISA are commonly used to detect seroconversion to flavivirus infections. However, faster, qualitative screening methods are needed for quicker diagnosis and better patient outcomes. Lateral flow assays (LFAs) can provide rapid results (5-15 mins) at the point-of-care, yet few commercial flavivirus antibody detection LFAs are available. We developed an LFA using novel chimeric viral antigens produced by genetically modifying the mosquito restricted Binjari virus (BinJV) to display the outer virion proteins of pathogenic viruses such as West Nile virus (WNV). The BinJV chimeric platform offers various advantages for diagnostic assay development, including rapid construction of new chimeras in response to emerging viral variants, safe, scalable antigen manufacturing, and structural indistinguishability to the wild-type pathogenic virion. As a demonstration of feasibility, we applied chimeric WNV (BinJV/WNV) antigen to LFA as the capture/test line reagent for detection of seroconversion of crocodilians to WNV – a virus affecting crocodilians on multiple continents. We verified the antigenic conservation of the chimera when applied to the LFA detection surface using monoclonal antibodies. Using well-characterised sera (n=60) from WNV seropositive or flavivirus naive Australian saltwater crocodiles (*Crocodylus porosus*), we illustrated 100% sensitivity and specificity, with results achieved in less than 15 minutes. The LFA further accurately detected seroconversion in animals experimentally infected with WNV. This qualitative screening method can be performed both inside and outside of a laboratory, and the assay design will guide the optimization of similar tests for vector borne virus infection detection in both humans and other animals.

## 1. Introduction

Flaviviruses such as West Nile (WNV), Zika (ZIKV), yellow fever (YFV), Murray Valley encephalitis (MVE) and Japanese encephalitis viruses (JEV) belong to a group of enveloped RNA viruses transmitted by arthropods that cause infections in humans and various vertebrate species (Simmonds et al. 2017). These viruses can have significant implications for global agriculture and import/export industries (Humblet et al. 2016; Mansfield et al. 2017). In the Australian context, the recent 2022 JEV range expansion caused by high rainfall, saw JEV rapidly spreading across the southern states, heavily affecting farmed and feral pig populations (Yakob et al. 2022). Similar conditions in previous years have also resulted in outbreaks of WNV in horses (Frost et al. 2012; Read et al. 2019). Another key industry affected by WNV, not only in Australia, but elsewhere in the world is the commercial crocodilian farming sector (Isberg et al. 2019; Jacobson et al. 2005; Roy et al. 2021). The Australian endemic strain of WNV, Kunjin (WNV_KUN_) currently causes substantial financial losses in the commercial crocodile-farming sector due to hide imperfections as a result of infections (Habarugira et al. 2020a; Isberg et al. 2019). Regular veterinary screening for flavivirus infections involves detecting genus or species-specific antibodies in blood using techniques such as enzyme-linked immunosorbent assays (ELISA), microsphere immunoassays (MIA), and immunofluorescence assays (IFA) (Hirota et al. 2013; Kerkhof et al. 2019). Confirmatory diagnosis relies on the plaque reduction neutralization test (PRNT), which detects virus-specific neutralizing antibodies and is considered the gold standard (Centers For Disease Control and Prevention (CDC) 2013). While these methods are sensitive, they pose challenges due to issues of specificity, cost, time consumption, and technical demands.

The global COVID-19 pandemic highlighted the utility of rapid antigen tests, such as lateral flow assays (LFAs), for point-of-care diagnostics (Peto et al. 2021). LFAs offer advantages such as low cost, rapid results, and minimal training requirements, making them an attractive alternative to traditional diagnostic methods (Ricks et al. 2021). Although there are limited LFAs available for WNV, companies like Response Biomedical Corporation and Vector Test Systems Inc. utilize antigen detection LFAs for mosquito screening and flaviviral antigen detection in mosquitoes and dead birds, respectively (Burkhalter et al. 2006; Pauvolid-Corrêa and Komar 2017). There are currently no LFA systems on the market for the specific detection of antibodies to WNV or many of the other flaviviruses that are responsible for disease in both a veterinary and human context.

Binjari virus (BinJV), an insect-specific flavivirus found in Australia, has demonstrated amenability with exchanging its structural prM and envelope protein genes with those of various vertebrate-infecting flaviviruses (Hobson-Peters et al. 2019). The resulting chimeric particles retain their insect-specific tropism and structural authenticity while not replicating in vertebrate cells (Hardy et al. 2021; Vet et al. 2020). These chimeric particles offer advantages such as high replication titres within mosquito cell lines and reduced bio-safety level requirements for their production and handling (Hobson-Peters et al. 2019). Currently, BinJV/WNV_KUN_ particles have been assessed as vaccine candidates (Habarugira et al. 2023; Vet et al. 2020) and applied to diagnostic assays such as ELISA and MIA (Hobson-Peters et al. 2019), but had not yet been used as an antigen in LFAs.

In this study, we aimed to evaluate an LFA format and develop a preliminary model capable of detecting flavivirus-specific antibodies using the BinJV/WNV_KUN_ chimera. This approach has the potential to enhance the capabilities of flavivirus surveillance and bridge gaps in diagnostic knowledge. To validate our approach, we chose saltwater crocodiles (*Crocodylus porosus*) as the target species given that they naturally acquire WNV infections (Isberg et al. 2019), develop robust immune responses against the pathogen (Habarugira et al. 2023; Habarugira et al. 2020a) and there are no rapid tests available for preliminary screening within the farm setting.

## 2. Experimental Section

### 2.1. Materials

Bovine serum albumin (BSA), sulfo-NHS (N-hydroxysulphosuccinimide), EDC (1-ethyl-3-(3-dimethylaminopropyl) carbodiimide hydrochloride), hydroxylamine, and Tween 20 were obtained from ThermoFisher Scientific. All additional reagents such as organic solvents, acids, salts, sugars, and alkalis were of analytical grade.

### 2.2. Design, construction, and operation of LFA

The final LFA device (Fig. 1A) consist of a 25 mm wide FF120HP (90-150 sec/4 cm) nitrocellulose membrane (Cytiva) with a 7 mm STD17 glass fibre conjugate pad (370 µm; Cytiva) impregnated with the dried gold conjugate. The opposite edge of the membrane was lapped by a 17 mm CF5 cotton linter absorbent pad (954 µm; Cytiva). The leading edge of the conjugate pad included a 17 mm Fusion 5 sample pad (370 µm; Cytiva). All components were attached to a backing card using a factory adhesive. Preliminary assessments were conducted by “spotting” 0.5 µL of test and control line reagents directly onto the nitrocellulose membrane. Membrane striping was carried out using a Biodot AD3060 membrane striping machine, and the strips were cut using a Kinematic Matrix 2600 guillotine. The test line was striped with BinJV/WNV_KUN_ while the assay control line consisted of a polyclonal antibody reactive to the gold conjugated antibody. The presentation of epitopes of the BinJV/WNV_KUN_ when applied to the membrane was confirmed using a goat anti-mouse Ig (Dako) conjugated gold nanoparticle (AuNP) and the control line comprised a polyclonal rabbit anti-goat Ig (Sigma). For the crocodile assay, goat, anti-alligator IgY (Novus) was conjugated to the AuNP, and the control line comprised a polyclonal rabbit anti-goat Ig (Sigma). Figure 1B demonstrates the LFA operational procedure, with 2.5 µL of sample applied, followed by 40-50 µL of running buffer (1xPBS, 0.5% T20). At 15 mins, the results were assessed visually and with the Leelu colorimetric reader (Lumos Diagnostics).

**Figure 1.**
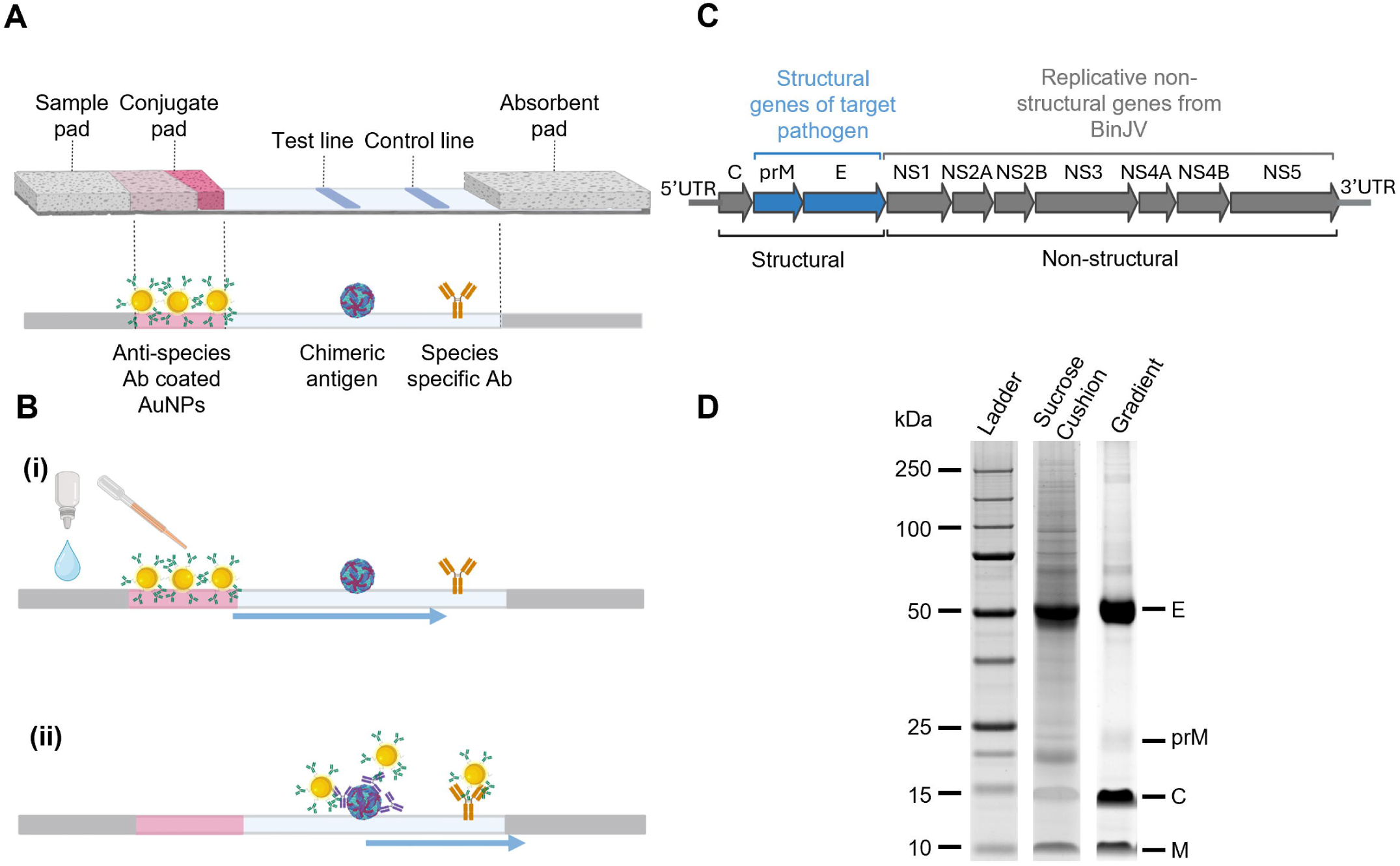
Design and operation of chimeric virus LFA. **(A)** Design of generic LFA incorporating chimeric BinJV chimeras. The design features goat anti-Ig conjugated AuNPs which can be interchanged depending on the target animal species, the BinJV chimera is applied to the test line and an anti-goat IgG is printed as the control line. **(B)** Illustration of chimeric virus LFA operation procedure. **(i)** 2 µL serum is applied and chased with 40-50 µL of running buffer, applied to the sample pad. **(ii)** The capillary flow of the membrane moves AuNPs from the conjugate pad to the test and control lines. In a positive sample, a brick red line is revealed due to the complexing of antibodies, conjugate, and chimeric virus on the test line. Unbound gold conjugate is trapped by the control line to designate successful running of the strip. **(C)** Schematic of the BinJV/WNVKUN chimera design. The WNVKUN prM/E genes are highlighted in blue and the BinJV capsid, non-structural protein genes and untranslated regions (UTRs) in grey. **(D)** SDS Page assessment of BinJV/WNVKUN virions purified by sucrose cushion only or by sucrose cushion followed by potassium tartrate gradient. Envelope protein **(E)** at 50 kDa, precursor to membrane (prM) at 20 kDa, capsid (C) at 15 kDa and membrane (M) at 10 kDa are indicated. **Created in part with Biorender.com**

### 2.3. Preparation of virus particles

Purified BinJV/WNV_KUN_ chimeric virions were applied as the test agent in the LFA. The chimeric virus was created by recombinantly inserting the prM/E structural protein genes of WNV_KUN_ in the backbone of BinJV (Hobson-Peters et al. 2019). To produce the chimeric viral antigen for application to LFA, BinJV/WNV_KUN_ was cultured and purified as previously described (Hobson-Peters et al. 2019). Briefly, C6/36 *Aedes albopictus* mosquito cells (ATCC CRL1660) grown in RPMI-1640 medium supplemented with 2% fetal bovine serum (FBS) were infected with BinJV/WNV_KUN_ at a multiplicity of infection (MOI) of 0.1. The culture supernatant containing released chimeric virus was harvested on days 3,5,7 and 10 post-infection, and the infected cell monolayer supplemented with fresh RPMI/2% FBS after each harvest. The virus supernatant was clarified by centrifugation at 3000 × RCF for 10 mins at 4 °C. The virus was precipitated overnight using 40% polyethylene glycol (PEG8000) and centrifuged at 8000 × RCF for 1 hour at 4 °C using an Avanti J-26 JLA10.5 rotor. The resulting virus pellet was resuspended in 1x NTE buffer (12 mM Tris at pH 8, 120 mM NaCl, 1 mM EDTA at pH 8) and ultracentrifuged at 28000 × RCF for 2 hours at 4 °C in Beckman Coulter SW32 tubes, through a 20% sucrose cushion. A sample of the sucrose cushion-purified virus was withheld for virion quantification analysis. For purity comparisons only, the precipitated virus pellet was resuspended in NTE buffer, clarified at 100 x RCF for 5 mins, and the resulting supernatant was loaded onto a 25-35% potassium tartrate gradient and centrifuged at 50,000 × RCF for 1 hour at 4 °C in Beckman Coulter SW60 tubes. The blue virus bands obtained from the gradient were harvested, buffer exchanged into NTE, and stored at 4°C.

The sucrose cushion and gradient purified viruses were analysed using SDS-PAGE as previously described, with the sucrose cushion concentration determined as detailed below (Choo et al. 2021). Briefly, the viral antigen samples were mixed with 4x LDS loading dye (ThermoFisher Scientific) and 1M DTT, heated at 90°C for 3 mins, and then subjected to gel electrophoresis (Hobson-Peters et al. 2019). To visualize the total protein, Sypro Ruby stain was used according to the manufacturer’s instructions. To quantify the purified virus, the intensity of the E protein monomer bands (50 kDa) was compared to BSA standards using ImageJ software.

### 2.4. Creation of gold conjugate

Antibodies were covalently conjugated to PEG Carboxyl-surfaced, 40 nm gold nanospheres (Bioready™ carboxyl gold nanoparticles, NanoComposix) following the manufacturer’s instructions. Briefly, the AuNPs were prepared for conjugation by applying EDC and sulfo-NHS stocks at a concentration of 10 mg/ml. Following incubation, the particles were pelleted, and the supernatant was removed. The particles were washed twice with K_2_HPO_4_ and PEG 20kDa before the addition of antibody (goat anti-mouse Ig or goat anti-alligator IgY) and incubation at RT on a rotator. After the incubation, a hydroxylamine quencher was added to stop the reaction, followed by an additional 10-minute incubation and a series of washes before storage in a diluent solution.

### 2.5. Assessment of anti-alligator IgY conjugate

To determine the suitability of the goat anti-alligator IgY conjugate for detection of saltwater crocodile antibodies, size exclusion chromatography purified crocodile IgY (Bizelli et al. 2015) was striped onto nitrocellulose membrane at a concentration of 1 mg/mL and dried at 37℃ for 1 hour. The anti-alligator IgY conjugate was applied to conjugate pads and dried. Strips were assembled as previously stated. To assess the binding capacity, 40-50 µL of running buffer (1xPBS, 0.5% T20) was applied to the sample pad. At 15 mins, the results were assessed visually and with the Leelu colorimetric reader.

### 2.6. Scanning electron microscopy of antigen applied to nitrocellulose membranes

BinJV/WNV_KUN_ spotted and unmodified nitrocellulose membranes were cut to size and mounted on adhesive carbon spots, dried and sputter coated to 5 nm with platinum. Imaging at resolutions ranging from 2-5 kilovolts (kv) was performed using a JEOL JSM-7001F Field Emission Scanning Electron Microscope (SEM) at the Centre for Microscopy and Microanalysis (CMM, The University of Queensland).

### 2.7. Antigen immobilisation and assessment

The optimal quantity of BinJV/WNV_KUN_ antigen for application to the LFA test was determined by applying a twofold serial dilution of purified virions as spots, starting at a concentration of 0.5 µg per spot, to pre-cut strips and dried. A characterised crocodile serum sample was applied to each of the strips followed by running buffer. Once run, strips were assessed by eye to determine variations of spot intensity variations between the different antigen dilutions.

For confirmation of epitope conservation, purified mouse, anti-WNV monoclonal antibodies (mAbs) against various domains of the viral envelope protein were applied as samples to an LFA. mAb binding to the BinJV/WNV_KUN_ test line was detected with anti-mouse Ig conjugated AuNPs. The mAbs were initially diluted to 1 mg/mL in sheep serum (as a sample matrix) and serially diluted 5-fold prior to application to the LFA. Each mAb was assessed in triplicate. The mAbs were BJ-6E6 (Harrison et al. 2020), 6B6C-1 (Roehrig et al. 1983) 3.67G (Adams et al. 1995) and 17D7 (Sanchez et al. 2005). As a negative isotype control, the mAb 7E3, which is reactive to negeviruses (Colmant et al. 2020), was also assessed.

### 2.8. Serum samples

Two sets of serum samples from *Crocodylus porosus* were employed in this project. One set was collected from commercial crocodile farms located in the greater Darwin region of the Northern Territory, Australia under UQ animal ethics permit SVS/354/20/NT Blood sample collection has been previously described (Habarugira et al. 2022).

A third set of sera originated from an experimental WNV infection study conducted in the Northern Territory under a protocol approved by the Charles Darwin University Animal Ethics Committee (Permit # A18004). The serum samples were heat inactivated at 56°C for 30 min before all diagnostic assay assessments.

The sera were initially screened in a pan-flavivirus blocking ELISA using 6B6C-1 as previously described (Habarugira et al. 2022). Those samples that returned a positive result in the 6B6C-1 blocking ELISA (percentage of inhibition >30%) were further assessed in a WNV-specific blocking ELISA using mAb 3.1112G (Blitvich et al. 2003a; Blitvich et al. 2003b; Hall et al. 1995) and virus neutralisation test using wild type WNV_KUN_ and established protocols at the Berrimah Veterinary Labs Northern Territory Australia (Uren 1993).

Sera from 10 animals enrolled in an experimental infection trial of *Crocodylus porosus* with WNV_KUN_ (Habarugira et al. 2020a) were randomly selected for assessment by LFA (control uninfected group n=3; 10^4^ IU WNV_KUN_ n=4, 10^5^ IU WNV_KUN_ n=3). For each crocodile in the WNV_KUN_ challenge groups, a pre challenge and/or day 1-6 post infection sample was assessed as a baseline negative sample as well as one or more samples taken at 1.5 to 3 months post WNV_KUN_ challenge. Samples from the uninfected crocodiles were taken at various timepoints during the trial. The sera had previously been assessed by the mAb 3.1112G WNV blocking ELISA described above and blocking ELISA positive samples confirmed in a WNV_KUN_ microneutralisation assay as previously described (Hall et al. 1995; Prow et al. 2014)

### 2.9. Strip stability assessment

To assess the LFA stability over time and under varying storage conditions, strips were tested in triplicate the day of manufacture, followed by subsequent testing at monthly intervals over a three-month period. Throughout the duration of the experiment, the LFA test strips were stored with desiccant, under three different temperature conditions: 4°C, room temperature, and 37 °C. The shelf life was calculated utilising the accelerated aging time (AAT) equation: 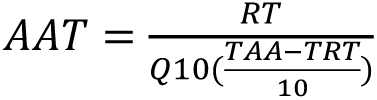 where RT = Real time, Q10 = ageing factor, TAA = accelerated aging temperature and TRT = ambient temperature (ASTM 2021; Porck 2000). Strip test line intensity results were measured and graphed with GraphPad Prism v.9-10 (GraphPad Software Inc.).

### 2.10. Data analysis

The colorimetric signals became visible with the naked eye within 5-10 mins and were then imaged using the Leelu reader (Lumos) at 15 mins. The reader measured three major metrics on each strip: Line area, background, and peak above background. This allowed for determination of test line strength as well as the differentiation between line indication and background “noise”. A lower limit of detection for the LFA was assigned as 0.05 (peak above background), which corresponded to the lowest visually (by eye) discernible indication of a line within the test region. Data exported from the reader as an excel file were plotted and compared to the previous serum assessments. For comparison with ELISA and PRNT results, peak above background readings were displayed graphically as multiplied by 10 or 100. All samples were assessed in triplicate. Virus neutralising titres were presented as the reciprocal value, whereby a value of 9 was the limit of detection. Data analysis and generation of graphs were performed using Microsoft Excel v.2312 (Microsoft) and GraphPad Prism v.9-10 (GraphPad Software Inc.).

## 3. Results

### 3.1. Production of chimeric virus particles and application to nitrocellulose membranes

Chimeric virions produced by the BinJV recombinant system have been assessed previously in ELISA and MIA diagnostic assays (Hobson-Peters et al. 2019). In the study described herein, we aimed to apply the chimeric particles to a paper-based assay for the serological detection of antibodies to WNV. The assay design comprised the application of BinJV/WNV_KUN_ chimera as the capture/test line and detection of sample derived anti-WNV antibodies with gold nanoparticles labelled with a suitable anti-species immunoglobulin (Fig 1A). A control line printed with a polyclonal antibody against the antibody conjugated AuNPs was used to ensure appropriate assay performance. Upon application of the sample and LFA running buffer, capillary flow facilitates the rehydration of the conjugated AuNPs and the migration of complexes along the membrane (Fig 1B). If antibodies reactive to the chimeric antigens were present in the sample, they bound to test line and complexed with the conjugate. Any unbound conjugate was captured on the control line antibody. The bound and unbound conjugate was indicated by the presence or absence of a red line on the test, indicating a positive or negative result respectively.

The structural and antigenic characterisation of the BinJV/WNV_KUN_ chimeric virions has been previously described (Hardy et al. 2021; Hobson-Peters et al. 2019). Its inability to replicate in vertebrate cells has also been confirmed (Hobson-Peters et al. 2019). In the current study, BinJV/WNV_KUN_ chimeric virus particles (Fig 1C), purified by sucrose cushion were assessed for application in the LFA. SDS-PAGE comparison of sucrose cushion-purified material against virions that had subsequently been taken through a potassium tartrate gradient showed that the preparations primarily consisted of mature virions, with an abundance of E protein at around 50kDa and negligible prM protein (Fig 1D). While it was clear that the gradient purification produced a cleaner product, overall, minimal host cell protein contaminants were observed following either purification method.

To determine the optimal amount of BinJV/WNV_KUN_ to be applied as the test line on the nitrocellulose membrane, a serial two-fold dilution of the antigen was performed, beginning at 0.5 µg/spot. The intensity of the test spot was determined using AuNPs conjugated with goat, anti-alligator IgY and a known WNV-positive crocodilian serum sample. Visual inspection revealed that antibody binding could be detected with as little as 0.03 µg of BinJV/WNV_KUN_ immobilised on the membrane, with the maximum spot intensity observed at 0.5 µg. Subsequent assays used an antigen concentration range between 0.8 and 1.2 mg/mL, corresponding to 0.4-0.6 µg per spot of virus.

Scanning electron microscopy (SEM) was employed to examine the application of the BinJV/WNV_KUN_ chimeric particles to the nitrocellulose membrane when spotted and dried. In comparison to the naked nitrocellulose (Fig 2A), the viruses were clearly visible, as spherical particles within the nitrocellulose matrix structure (Fig 2B-D). The imaging indicated that the chimeric particles were dispersed evenly with no aggregation evident. The apparent spherical appearance of the particles suggested that structural integrity was maintained when applied to LFA.

**Figure 2.**
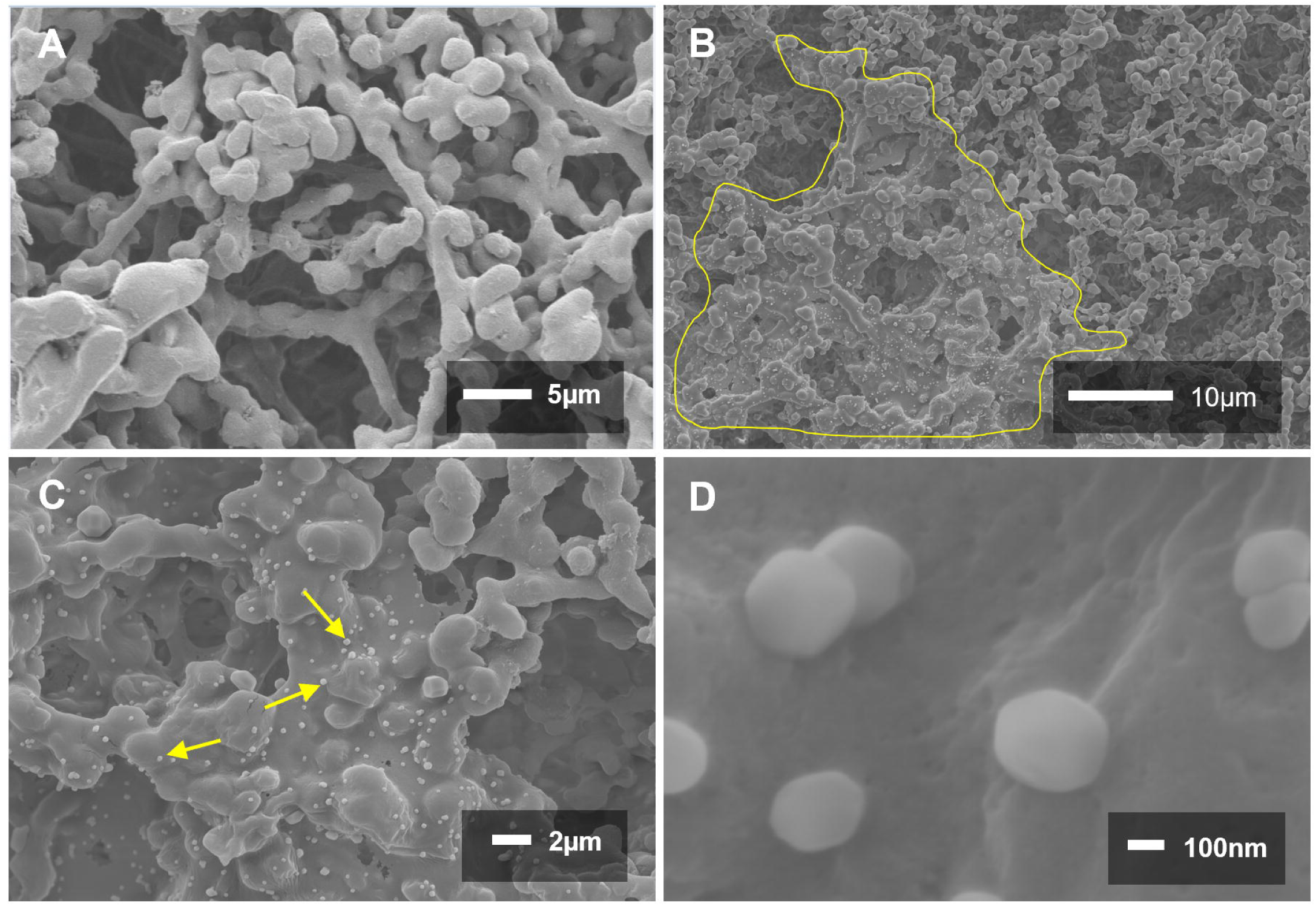
BinJV/WNVKUN applied to nitrocellulose. **(A)** SEM of naked nitrocellulose membrane at 3000x magnification. **(B)** SEM of BinJV/WNVKUN spotted on nitrocellulose membrane. Region surrounded by yellow border indicates dried, immobilised viral particles in striping fluid. **(C-D)** SEM of BinJV/WNVKUN spotted on nitrocellulose membrane. Representative chimeric particles are indicated by yellow arrows. Imaging performed at 1400x **(B),** 5000x **(C)** and 95000x **(D)** magnification.

### 3.2. Epitope presentation assessment

To ensure the appropriate display of antigenic epitopes when the BinJV/WNV_KUN_ chimera was immobilised on the nitrocellulose membrane, a panel of mAbs targeting different E protein domains of WNV_KUN_ were assessed in LFA and detected with goat, anti-mouse Ig conjugated AuNPs (Fig 3A). All anti-WNV mAbs bound strongly in the system, returning a positive result even at low concentrations, confirming appropriate presentation of the epitopes (Fig 3B). There was no reactivity with an isotype control mAb.

**Figure 3.**
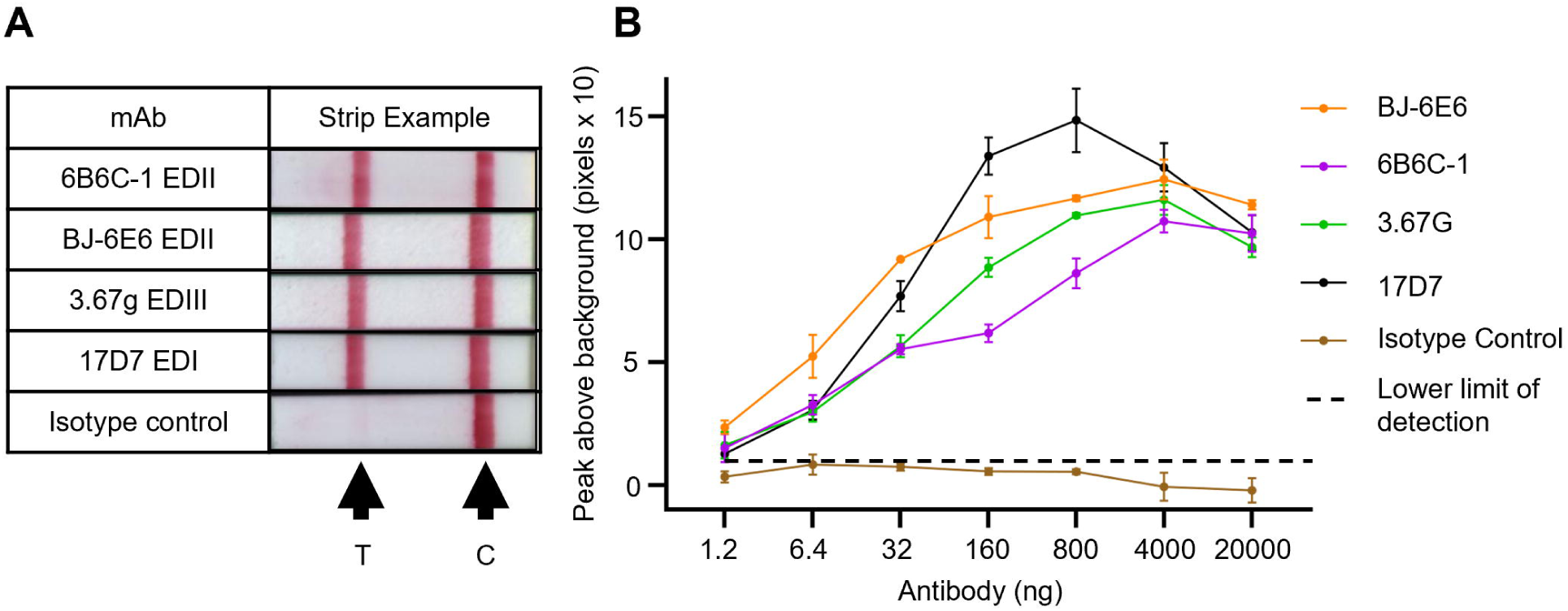
Epitope presentation assessment. Epitope display assessment was performed using an LFA system of BinJV/WNVKUN immobilised as the test line, goat anti mouse IgG conjugated AuNPs and murine mAbs as the test sample. **(A)** Representative LFA peak images for each mAb (800 ng). T= test line and C = control line. **(B)** Results of five-fold serial dilution of mAbs tested in LFA in triplicate. The test line peak above background, as determined by the Leelu reader and multiplied by 10, was plotted. mAbs assessed: Envelope Domain 1 (EDI) (17D7), EDII (BJ-6E6, and 6B6C-1), and EDIII (3.67G). mAb (7E3) was utilised as an isotype control (negative).

### 3.3. Application of the BinJV/WNV_KUN_ LFA for assessment of WNV-immune sera

Crocodilians represent an ideal test case for the assessment of the BinJV/WNV_KUN_ LFA. Australian saltwater crocodiles become infected with WNV, developing skin lesions known as “pix” because of the infection (Habarugira et al. 2020a; Isberg et al. 2019), and thus monitoring farmed animals for exposure to the virus is desirable. Due to the absence of commercially accessible anti-crocodile IgY antibodies, a goat, anti-alligator IgY polyclonal antibody (pAb) was employed in the LFA system for conjugation to the AuNPs.

To confirm sensitive detection of crocodile IgY by the anti-alligator IgY conjugate, purified crocodile IgY was applied to the nitrocellulose prior to probing with the anti-alligator IgY conjugate. A strong positive result was detected indicating suitability for the use of the anti-alligator conjugate to detect crocodile IgY (Fig S1). A preliminary assessment of both sucrose cushion and gradient purified BinJV/WNV_KUN_ was conducted to identify the degree of purification required for the assay, with the much less laborious sucrose cushion method yielding adequate results to continue. The quantity of antigen required for test line was determined by eye as 0.5 – 1 mg/mL (Fig 4A). Confirmatory function of the LFA was assessed using one sample from a WNV-naïve crocodile and samples from three WNV-seropositive crocodiles, as confirmed by VNT. The test line intensity ranged from visibly weak to strong and an absence of a visible test line for the sero-negative sample, indicating that the set-up was sound (Fig 4B).

**Figure 4.**
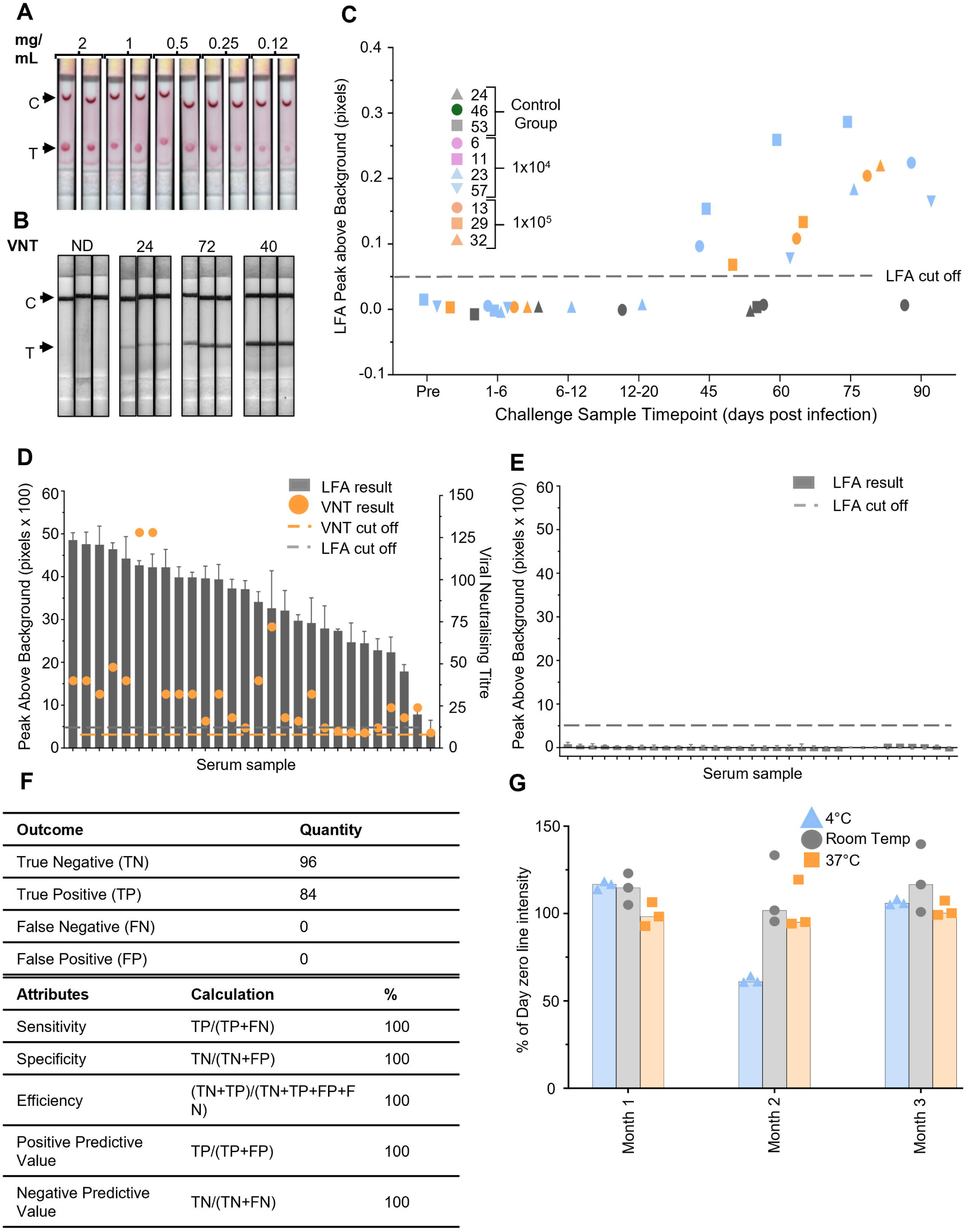
Crocodile serum analysis by LFA. **(A)** Two-fold serial dilutions (2, 1, 0.5, 0.25 and 0.12 mg/mL) of BinJV/WNVKUN in PBS were applied to nitrocellulose strips, probed with WNV-positive crocodile serum, and detected with anti-alligator IgY-conjugated AuNPs. T = test, C = control. **(B)** Strip examples showing a negative, low, medium, and high test line signal on crocodile serum assessment. The corresponding WNV VNT value is provided above the triplicate strips. T = test line and C = control line. Panels of crocodile sera as assessed by LFA. **(C)** Assessment of animals in an experimental infection trial with WNVKUN. Control group (grey) and infected groups (blue – 1x104 IU WNVKUN, orange – 1x105 IU WNVKUN), sampled pre-challenge, 1-20 days, 5.5 months, 6 months, 6.5 months, and 7 months post challenge. **(D)** 28 WNV virus neutralisation test (VNT) and WNV blocking ELISA positive crocodile serum samples; **(E)** 32 flavivirus negative samples. Samples were assessed by LFA in triplicate, strips imaged and analysed using Leelu reader (left y axis, grey column), and compared to VNT results (right axis, orange spot). Grey broken line indicates pre-defined lower limit cut off for LFA. Broken orange line indicates pre-defined lower limit cut off for VNT. **(F)** Number of strips assessed as true positive, true negative, false positive and false, negative based on data from panels D and E. **(G)** Stability assessment of LFA at 4℃, room temperature and 37℃ over a 3-month period. Strips were stored with desiccant at each temperature and assessed in triplicate at each time point. Results graphed as a percentage of day zero test line intensity. Strips were tested using one WNV positive and one negative crocodile serum sample.

The final assay design for testing of crocodiles for seroconversion to WNV consisted of a test line striped with 1 mg/mL BinJV/WNV_KUN_ and a control line striped with rabbit anti-goat Ig to detect any unbound goat anti-alligator IgY conjugate. Confirmation of detection of WNV seroconversion in crocodiles by the LFA was provided through assessment using a panel of experimentally infected crocodiles previously described (Habarugira et al. 2020a). Sera from 10 animals of an uninfected group (n=3), infected at 10^4^ IU WNV_KUN_ (n=4) and 10^5^ IU WNV_KUN_ (n=3) were assessed (Fig 4C) Each sample was tested in triplicate and strips were read with the Leelu colorimetric reader. An assay cutoff to discriminate between visually negative and positive samples was defined as 0.05 (pixels) peak above background. Samples falling below that value are considered negative and those above are positive. As previously stated, the cutoff values were determined by subjective visual assessment. Samples taken pre-challenge or within one to six days post challenge for the 10^4^ IU and 10^5^ IU WNV_KUN_ animals returned negative LFA results, consistent with the results of the blocking ELISA (Fig 4C, Table S1). All post-challenge serum samples taken from 1.5 months onwards were positive by LFA, typically with increasing test line intensity for those animals sampled numerous times between the 1.5 to 3 months post-challenge and consistent with rising VNT results (Fig 4C, Table S1). For all samples assessed from crocodiles in the control unchallenged group, the LFA assessments returned the expected negative result over the duration of the trial. Together, these data indicated that the LFA could accurately detect seroconversion to WNV 1.5 months post infection.

A larger panel of samples collected from crocodiles housed in a farm setting was subsequently assessed. The 60 crocodile serum samples, separated into two panels consisting of 28 anti-WNV positive samples (Table S2) and 32 antibody negative samples (Table S3).

The serological status of each crocodile sample used in the confirmatory panels was initially determined by the flavivirus-generic 6B6C-1 blocking ELISA (Habarugira et al. 2022). Samples that returned a positive result (percentage of inhibition >30%) were further assessed in the WNV specific blocking ELISA and WNV_KUN_ virus neutralisation test (VNT) (Tables S2, S3). When assessed in LFA, all samples confirmed as seropositive for WNV antibodies returned a visually positive result in LFA (Fig 4D) of varying intensity, but with peak reading measurements above the threshold defined to demarcate a positive or negative sample. Of note, however, is that there was no correlation between the VNT titre and the intensity of the LFA test line. Samples confirmed negative in the flavivirus blocking ELISA did not yield an observable line and when assessed by the Leelu reader, the peak readings consistently remained below the predetermined cutoff point (Fig 4E).

Based on the findings from both confirmatory panels, the LFA exhibited 100% sensitivity in identifying anti-WNV antibodies within the crocodile serum (Fig 4F). No false positives were detected when testing flavivirus antibody-negative samples, providing a specificity of 100% (Fig 4F).

### 3.4. Assay shelf-life

LFA strips, prepared as described above, were evaluated using one positive and one negative crocodile serum sample after storage at temperatures of 4°C, room temperature (RT), and 37°C spanning three months. The strips were evaluated in triplicate on the day of preparation (day zero), and monthly intervals thereafter. Test line intensity was documented at each time point and compared to the line intensity at day 0 (Fig 4G). The positive sample remained clearly positive at each time point of strip storage with minimal variation of intensity of test line. Similarly, upon testing the negative sample, no false positive results were observed. The strips displayed consistency across the timepoints with a slight reduction in performance in month two for the strips stored at 4°C, however month three saw the 4°C strips return to 100% of day zero. Extrapolating the potential shelf life using the Arrhenius test (ASTM 2021; Porck 2000) based on the strips held at 37°C, the shelf life was determined to be 240 days when held at room temperature.

## 4. Discussion

LFAs, particularly when deployed in field settings, prove to be highly advantageous as screening tools, rapidly identifying samples warranting further testing within 20 mins (Nan et al. 2023). With their quick turnaround time, LFAs enable high throughput testing without prolonged incubation periods or waiting times, offering an attractive alternative to conventional screening methods. However, few LFAs able to detect seroconversion to flaviviral infections in the medical or veterinary fields (De Filette et al. 2012; Dhanze et al. 2020) have been reported, with even fewer taken through to commercialisation (Habarugira et al. 2020b). Herein we paired the BinJV chimeric technology with LFA to produce a system that offers safe, rapid production of antigen coupled with a quick, simple, and sensitive way to diagnose WNV infections in crocodiles.

As a test case, we chose to develop an assay against WNV. WNV has a global spread that has seen human outbreaks in numerous countries including Romania, Russia, Israel, Egypt, India, France, South Africa, Canada, U.S and Mexico ranging from mild febrile illness to neurological infections and encephalitis (Chancey et al. 2015). Avians, horses and crocodilians are also susceptible to WNV infections with outbreaks occurring as widespread as human infections. Notably, the endemic strain of WNV (Kunjin) regularly cause sporadic equine infections across northern Australia with an increase of cases as recently as 2011 due to unusually wet weather providing optimal mosquito breeding conditions (Frost et al. 2012; Prow et al. 2016). Like human infections, horses can suffer neurological disease, leading to fatal outcomes. Crocodiles are also susceptible to infections; however, in the contest of WNV_KUN_, the effects are often less dramatic and only lead to the development of small lesions in the hide called pix which result in financial damage to the farming industry (Habarugira et al. 2020a). In other crocodilian species, however, infection with other strains of WNV can cause severe clinical signs, often resulting in death, as is the case with alligators infected with the North American strains of WNV in alligators (Jacobson et al. 2005; Klenk et al. 2004) It is important to note that due to the similarity of crocodilians in general, the LFA developed herein may be employed for other species of crocodilians such as alligator and cayman, providing a globally employable screening tool (Habarugira et al. 2022).

Currently there is a worldwide gap in the testing capabilities for the detection of WNV infections worldwide in crocodiles, with testing currently reliant on laboratory-based assays such as PRNT, blocking ELISA or qRT-PCR (Habarugira et al. 2020b; Hirota et al. 2013). In many crocodilian farming nations, laboratory resources are not available (Tosun 2013), making the utility of field based LFAs highly attractive. Concerns regarding sensitivity and specificity of LFAs have led to a void not only in the knowledge, but also the commercial sphere (Varghese et al. 2023). The employment of the “plug-and-play” BinJV platform technology within LFA lends itself to the capacity for modifications, not only for the detection of antibodies to different flaviviruses, but also for use in screening different animal species. A rapid screen that can be performed pen-side at transition stages within a farm offers a quick solution for screening populations, allowing for management changes to reduce mosquito-independent transmission within the farm (Habarugira et al. 2020a).

The chimeric BinJV virus particles have demonstrated their utility in other diagnostic platforms such as ELISA and MIA, performing similarly to that of the wild-type counterpart (Harrison et al. 2020; Hobson-Peters et al. 2019). Both the wild-type virus and BinJV/WNV_KUN_ have been assessed head-to-head in comprehensive structural analysis, proving indistinguishable from each other (Hardy et al. 2021; Hobson-Peters et al. 2019). We saw that application of whole chimeric virions based on the BinJV platform in LFAs facilitated presentation of a complex antigenic structure likely mirroring the actual virus encountered during infection. The presence of multiple antigenic sites on BinJV/WNV_KUN_ may increase sensitivity, reducing the likelihood of false negatives compared to an assay employing a single subunit protein or peptide as the capture antigen. In this study we applied the chimeric particles to nitrocellulose membranes and assessed the availability of each of the surface-presented epitopes by probing with specific mAbs. This, coupled with the SEM imaging suggested that chimeric particle integrity had been maintained throughout the LFA application process.

Testing to determine the optimal quantity of chimeric virions as the test reagent, yielded a plateau of test dot intensity when 0.5 mg/mL to 2 mg/mL dots of the chimeric antigen were applied and likely explained by the principle of antigen excess. That is, the ratio of antibody binding sites within the test-dot-applied chimeric virions far exceeded that of the binding antibodies in the sample (Jacobs et al. 2015). A final concentration of 0.8-1.2mg/mL chimeric antigen was settled upon to avoid the likelihood of a hook effect which could cause false negative results should the antigen concentration become too high (Gillet et al. 2009; Strait 2006).

To challenge the accuracy of the LFA set-up for the detection of seroconversion to WNV infection in crocodiles, serum taken from 3-4 animals in three different groups of a WNV_KUN_ experimental challenge trial were analysed (Habarugira et al. 2020a). Analysis of the serum from uninfected animals revealed a negative LFA result, regardless of when the sample was taken during the trial and was concordant with the negative result obtained in the blocking ELISA. However, specific seroconversion to WNV 1.5 months onwards for those animals that were experimentally infected with WNV_KUN_ typically with an increasing line intensity as the trial continued and consisted with increasing VNT values. For the experimentally challenged animals, pre-challenge, or days 1-6 post challenge samples served as a baseline negative control and returned a negative LFA result. Days 1-6 post challenge represent the key time frame for viraemia, as demonstrated by the detection of WNV_KUN_ RNA in sera of these cohorts (Habarugira et al. 2020a). In these animals, detectable levels of neutralising antibody to WNV_KUN_ was not evident until day 16 at the earliest, with most animals having detectable seroconversion by day 21. Further examination of samples collected from day 6 post infection onwards would provide a more precise indication of how early the LFA could detect seroconversion to WNV_KUN_.

The larger, confirmatory panels of crocodile serum further substantiated the LFAs capacity to accurately detect seroconversion to WNV_KUN_, resulting in 100% for both sensitivity and specificity across all samples tested. These serum samples were taken from animals held under standard farming conditions, indicating the utility of the LFA to assess for seroconversion in a farm setting.

Some of the samples were a subset of a larger group of 500 samples previously tested for various flaviviral antibodies (Habarugira et al. 2022). Of these, 68.5% were found negative for flavivirus using blocking ELISA. Screening such a large number of samples with blocking ELISA techniques not only demands highly skilled operators but also entails significant financial and time investments. The LFA described could reduce testing time to <20 mins.

The stability testing of the strips showed that the temperature ranges tested (4℃, RT and 37℃) had little impact on the results, indicating that over the testing period (three months), stability was maintained. The accelerated aging time equation provided a prediction of long-term storage capacity based on the real time experiment conducted over three months. While this prediction provides preliminary information about the longevity of the assay, further testing over a greater temperature range and a longer timeframe would provide more detailed information as to the long-term storage capability of the LFA as well as direction for optimum storage conditions.

With a simple replacement of the anti-alligator IgY conjugate with that of another animal, other at-risk animals such as horses, birds or humans could be screened for WNV exposure. Likewise, the replacement of the BinJV/WNV_KUN_ test line reagent with another BinJV chimera such as BinJV/JEV or BinJV/Murray Valley encephalitis, would allow screening for infections in at risk populations.

The employment of the described LFA for screening large populations would be invaluable due to its simplicity, low cost, and rapid throughput, providing results in under 20 mins. While this study employed a colorimetric reader to obtain data that could be benchmarked against established methods, the practical employment of the LFA “pen side” would allow for visual determination by eye to further reduce setup costs and hands on time. The practical utility of this LFA extends beyond financial and temporal benefits. Given the transmission dynamics of WNV, rapid screening of crocodile populations could mitigate risks associated with mosquito-independent transmission via the fecal-oral route, thereby safeguarding farm profitability. Additionally, screening for WNV exposure in juvenile crocodiles can inform vaccination strategies, ensuring more effective vaccination regimes by accounting for maternal antibodies.

## 5. Conclusion

This study validated the BinJV/WNV_KUN_ LFA as a robust method for detecting flavivirus-reactive antibodies within crocodile serum samples. The use of an anti-alligator IgY conjugate, sample categorisation based on VNT and blocking ELISA results, and the establishment of a reliable assay cutoff contributed to the reliability of the study’s findings. We believe that this assay will form a blueprint for application to other species that become infected with WNV such as birds, horses and humans (Habarugira et al. 2020b; Root and Bosco-Lauth 2019). The potential application of BinJV chimeric antigens in various LFA formats holds promise for detecting a range of mosquito-borne viral infections. This qualitative screening approach has implications for both laboratory and field settings, serving as a guide platform to develop similar tests targeting different viral infections in humans and other animals.

## Supplementary Materials

Table S1: The serological status of WNVKUN experimentally infected crocodile samples taken on days post infection, determined by VNT, Blocking ELISA and then assessed by LFA. Table S2: The serological status of each crocodile sample determined by VNT, flavivirus-generic 6B6C-1 blocking ELISA, WNV specific 3.1112G blocking ELISA and then assessed by LFA. Table S3: The serological status of each crocodile sample determined by flavivirus-generic 6B6C-1 blocking ELISA and then assessed by LFA. Figure S1: Anti-alligator conjugate compatibility assessment.

## Author contributions

Conceptualisation, J.H-P., R.A.H, J.Mac. and R.A.J., methodology, R.A.J, G.H., J.H-P. R.A.H., S.R.I. J.J.H., C.B.H and C.S.H, validation, J.H-P, R.A.J. and G.H., formal analysis, R.A.J., G.H., J.H-P., H.B-O., R.A.H., and S.R.I., investigation, R.A.J., J.H-P., J.J.H., and M.M., resources, R.A.J., G.H., J.H-P., J.J.H., J.M., S.R.I., J.Mac., M.M., S.S.D., L.M., H.B-O. and R.A.H., data curation, R.A.J., G.H., and J.H-P., writing – original draft preparation, R.A.J., J.H-P., and R.A.H., writing – review and editing, all authors, visualisation, R.A.J. and J.H-P., supervision, J.H-P., R.A.H., C.B.H. and C.S.H. project administration, J.H-P., S.R.I. and R.A.H, funding acquisition, J.H-P., S.R.I, H.B-O., C.B.H., and R.A.H. All authors have read and agreed to the published version of the manuscript.

## Institutional review board statement

This study was conducted in accordance with the University of Queensland research ethical guidelines. Ethical approval for this study was obtained from the University of Queensland’s Native and Exotic Wildlife and Marine Animal Ethics Committee (SVS/354/20/NT) and from the Charles Darwin University Animal Ethics Committee (Permit # A18004). The Centre for Crocodile Research has a license to conduct research on animals with the animal welfare unit of the Northern Territory government (permit no. 061). During sample collection, animal care and use protocols adhered to the Animal Welfare Regulations 2000 of the Northern Territory of Australia and Code of Practice on the Humane Treatment of Wild and Farmed Australian Crocodiles.

## Informed consent statement

Not applicable.

## Acknowledgments

The authors would like to acknowledge the Crocodile Farm staff and Berrimah veterinary labs management and staff for providing access to crocodile serum samples, as none of the subsequent vaccine or diagnostic work could be conducted without their painstaking work. We acknowledge the facilities and the scientific and technical assistance of the Australian Microscopy and Microanalysis Research Facility at the Centre for Microscopy and Microanalysis, University of Queensland. Finally, we acknowledge BioCifer Pty Ltd for the use of LFA manufacturing equipment.

