## Supplementry files for "Application of chimeric antigens to paper-based diagnostics for detection of West Nile virus infections of *Crocodylus porosus –* a novel animal test case"

### Supplementary data

***Table S1. Assessment of $WNV_{KUN}$ experimentally infected crocodile samples by LFA.***

**$WNV_{KUN}$ experimentally infected crocodile samples**

| ID | DPI* | $WNV_{KUN}$ VNT Titre and Classification** | Blocking ELISA % Inhibition^ | LFA Mean Peak Above Background^^ | LFA Pos/Neg§ | Strip Image |
|---|---|---|---|---|---|---|
| 6 | 6 | <20 Negative | <30 | 0.0054 | Neg | |
| | 45 | 20 Positve | <30 | 0.0966 | Pos | |
| | 90 | 320 Positive | 84.4 | 0.2239 | Pos | |
| 11 | 0 | <20 Negative | 44.0 | 0.0149 | Neg | |
| | 4 | <20 Negative | <30 | -0.0018 | Neg | |
| | 45 | 80 Positive | <30 | 0.1536 | Pos | |
| | 60 | 80 Positive | 78.9 | 0.2589 | Pos | |
| | 75 | 320 Positive | 77.3 | 0.2864 | Pos | |
| 13 | 5 | <20 Negative | <30 | 0.0035 | Neg | |
| | 6 | <20 Negative | <30 | 0.0049 | Neg | |
| | 60 | 40 Positive | 49.4 | 0.1082 | Pos | |
| | 75 | 160 Positive | 63.1 | 0.2041 | Pos | |
| 23 | 6 | <20 Negative | <30 | -0.00385 | Neg | |
| | 9 | <20 Negative | 80.0 | 0.00405 | Neg | |
| | 15 | <20 Negative | <30 | 0.00795 | Neg | |
| | 75 | 640 Positive | 84.0 | 0.1838 | Pos | |
| 24 | 16 | <20 Negative | 33.5 | 0.00435 | Neg | |
| | 60 | <20 Negative | <30 | -0.0021 | Neg | |
| 29 | 0 | <20 Negative | <30 | 0.00295 | Neg | |
| | 45 | <20 Negative | <30 | 0.06815 | Pos | |
| | 60 | 20 Positive | 51.5 | 0.1333 | Pos | |
| 32 | 1 | <20 Negative | <30 | 0.0038 | Neg | |
| | 75 | 160 Positive | 82.7 | 0.2202 | Pos | |
| 46 | 1 | <20 Negative | <30 | -0.0076 | Neg | |
| | 60 | <20 Negative | <30 | 0.00345 | Neg | |
| 53 | 17 | <20 Negative | <30 | -0.0008 | Neg | |
| | 20 | <20 Negative | <30 | -0.00495 | Neg | |
| | 60 | <20 Negative | <30 | 0.00695 | Neg | |
| | 90 | <20 Negative | <30 | 0.0063 | Neg | |
| 57 | 0 | <20 Negative | <30 | 0.00435 | Neg | |
| | 3 | <20 Negative | <30 | 0.00205 | Neg | |
| | 60 | 20 Positive | 44.8 | 0.07785 | Pos | |
| | 90 | 2560 Positive | 34.4 | 0.1647 | Pos | |

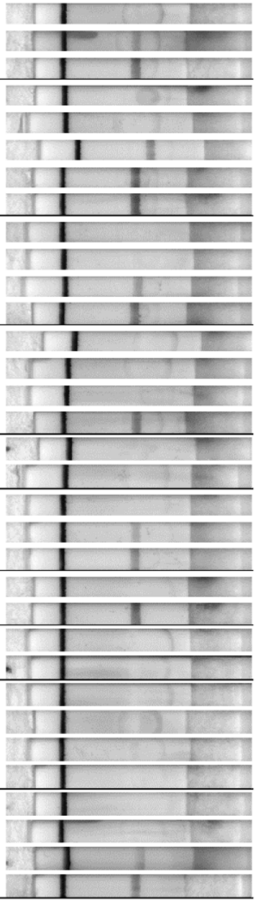

*Days post infection, **VNT titre cutoff 20, ^Blocking ELISA % cutoff 30, ^^Background subtracted from peak reading of LFA test line using Leelu reader, §LFA positive/negative based on a peak above background cutoff of 0.05.

*Table S2. Determination of seropositive samples and LFA results.*

**Positive crocodile serum samples**

| ID | $WNV_{KUN}$ VNT Titre and Classification* | Blocking ELISA % Inhibition Flavi** | KUNV^ | Conclusion | LFA Mean Peak Above Background^^ | Strip Example |
|---|---|---|---|---|---|---|
| 151586 | 9 Trace | 95 | 87 | KUNV antibody positive | 0.068 | 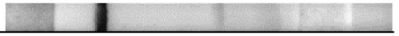 |
| 151587 | 24 Postive | 100 | 95 | KUNV antibody positive | 0.223 | 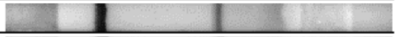 |
| 151589 | 12 Positive | 100 | 100 | KUNV antibody positive | 0.228 | 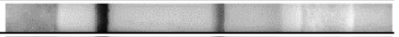 |
| 151590 | 40 Positive | 91 | 100 | KUNV antibody positive | 0.443 | 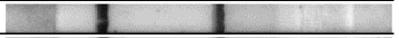 |
| 151592 | 32 Positive | 95 | 100 | KUNV antibody positive | 0.398 | 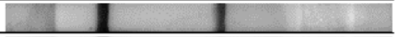 |
| 151594 | 24 Positive | 86 | 100 | KUNV antibody positive | 0.078 | 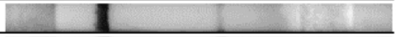 |
| 151595 | 72 Positive | 99 | 100 | KUNV antibody positive | 0.326 | 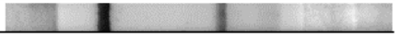 |
| 151600 | 128 Positive | 100 | 100 | KUNV antibody positive | 0.422 | 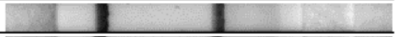 |
| 151601 | 128 Positive | 88 | 100 | KUNV antibody positive | 0.426 | 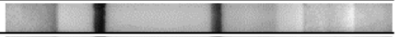 |
| 151604 | 40 Positive | 100 | 100 | KUNV antibody positive | 0.486 | 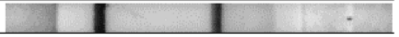 |
| 151605 | 48 Positive | 100 | 100 | KUNV antibody positive | 0.464 | 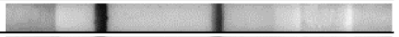 |
| 151606 | 40 Positive | 100 | 100 | KUNV antibody positive | 0.476 | 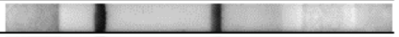 |
| 152580 | 9 Trace | 100 | 100 | KUNV antibody positive | 0.244 | 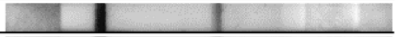 |
| 152581 | 16 Positive | 86 | 100 | KUNV antibody positive | 0.298 | 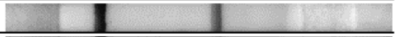 |
| 152582 | 18 Positive | 82 | 100 | KUNV antibody positive | 0.321 | 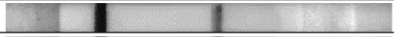 |
| 152583 | 16 Positive | 100 | 100 | KUNV antibody positive | 0.396 | 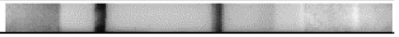 |
| 152585 | 32 Positive | 100 | 100 | KUNV antibody positive | 0.394 | 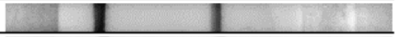 |
| 152589 | 18 Positive | 100 | 100 | KUNV antibody positive | 0.373 | 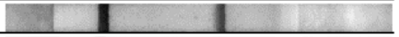 |
| 152590 | 18 Positive | 83 | 99 | KUNV antibody positive | 0.179 | 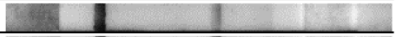 |
| 152591 | 32 Positive | 100 | 100 | KUNV antibody positive | 0.399 | 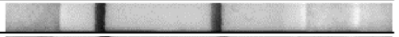 |
| 152593 | 9 Trace | 97 | 100 | KUNV antibody positive | 0.247 | 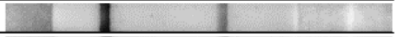 |
| 152594 | 32 Positive | 97 | 100 | KUNV antibody positive | 0.292 | 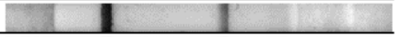 |
| 152596 | 32 Positive | 100 | 100 | KUNV antibody positive | 0.422 | 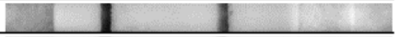 |
| 152597 | 32 Positive | 89 | 100 | KUNV antibody positive | 0.475 | 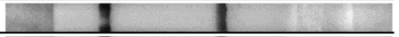 |
| 152600 | 12 Positive | 99 | 100 | KUNV antibody positive | 0.371 | 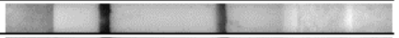 |
| 152601 | 10 Positive | 98 | 100 | KUNV antibody positive | 0.273 | 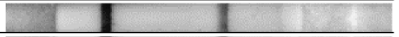 |
| 152602 | 12 Positive | 100 | 100 | KUNV antibody positive | 0.279 | 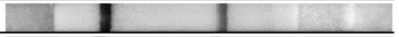 |
| 152603 | 40 Positive | 100 | 100 | KUNV antibody positive | 0.341 | 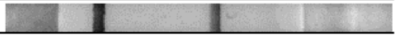 |

*VNT titre cutoff 10, **6B6C-1 Pan-flavivirus blocking ELISA % cutoff 30, ^3.1112G WNV specific blocking ELISA % cutoff 30, ^^Background subtracted from peak reading of LFA test line using Leelu reader Cutoff 0.05.

*Table S3. Determination of seronegative samples and LFA results.*

**Negative crocodile serum samples**

| ID | Blocking ELISA % Inhibition Flavi* | Conclusion | LFA Mean Peak Above Background** | Strip Example |
|---|---|---|---|---|
| A446 | <30 | Flavi antibody negative | 0.0076 | 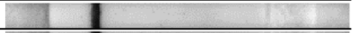 |
| A448 | <30 | Flavi antibody negative | 0.0051 | 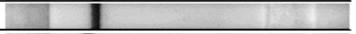 |
| A487 | <30 | Flavi antibody negative | 0.0047 | 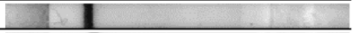 |
| A445 | <30 | Flavi antibody negative | 0.0046 | 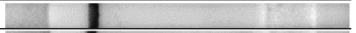 |
| 151585 | <30 | Flavi antibody negative | 0.0038 | 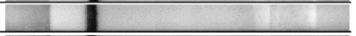 |
| A491 | <30 | Flavi antibody negative | 0.0032 | 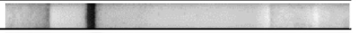 |
| A449 | <30 | Flavi antibody negative | 0.0031 | 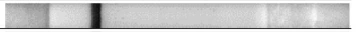 |
| A444 | <30 | Flavi antibody negative | 0.0031 | 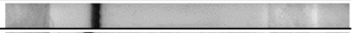 |
| A499 | <30 | Flavi antibody negative | 0.0027 | 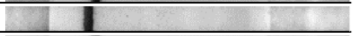 |
| A447 | <30 | Flavi antibody negative | 0.0025 | 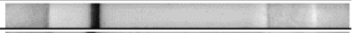 |
| A452 | <30 | Flavi antibody negative | 0.0024 | 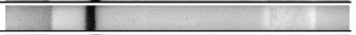 |
| A450 | <30 | Flavi antibody negative | 0.0021 | 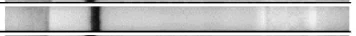 |
| A456 | <30 | Flavi antibody negative | 0.0020 | 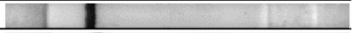 |
| A442 | <30 | Flavi antibody negative | 0.0019 | 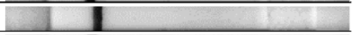 |
| A495 | <30 | Flavi antibody negative | 0.0019 | 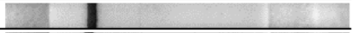 |
| A500 | <30 | Flavi antibody negative | 0.0017 | 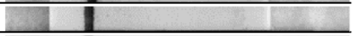 |
| A453 | <30 | Flavi antibody negative | 0.0014 | 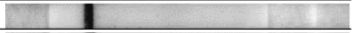 |
| A494 | <30 | Flavi antibody negative | 0.0014 | 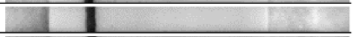 |
| A489 | <30 | Flavi antibody negative | 0.0012 | 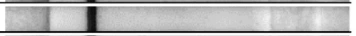 |
| A455 | <30 | Flavi antibody negative | 0.0010 | 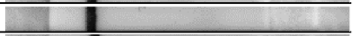 |
| A451 | <30 | Flavi antibody negative | 0.0009 | 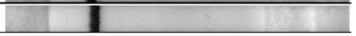 |
| A497 | <30 | Flavi antibody negative | 0.0006 | 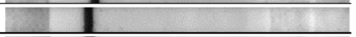 |
| A498 | <30 | Flavi antibody negative | 0.0005 | 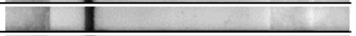 |
| A492 | <30 | Flavi antibody negative | 0.0004 | 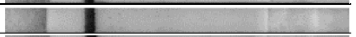 |
| 151584 | <30 | Flavi antibody negative | 0.0003 | 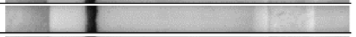 |
| A454 | <30 | Flavi antibody negative | 0.0001 | 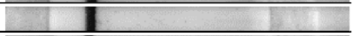 |
| A443 | <30 | Flavi antibody negative | 0.0000 | 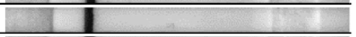 |
| A490 | <30 | Flavi antibody negative | 0.0000 | 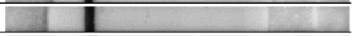 |
| A496 | <30 | Flavi antibody negative | -0.0001 | 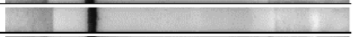 |
| A441 | <30 | Flavi antibody negative | -0.0001 | 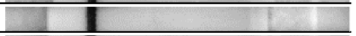 |
| A493 | <30 | Flavi antibody negative | -0.0005 | 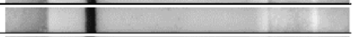 |
| A488 | <30 | Flavi antibody negative | -0.0015 | 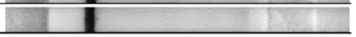 |

*6B6C-1 Pan-flavivirus blocking ELISA % cutoff 30, ** Background subtracted from peak reading of LFA test line using Leelu reader. Cutoff 0.05.

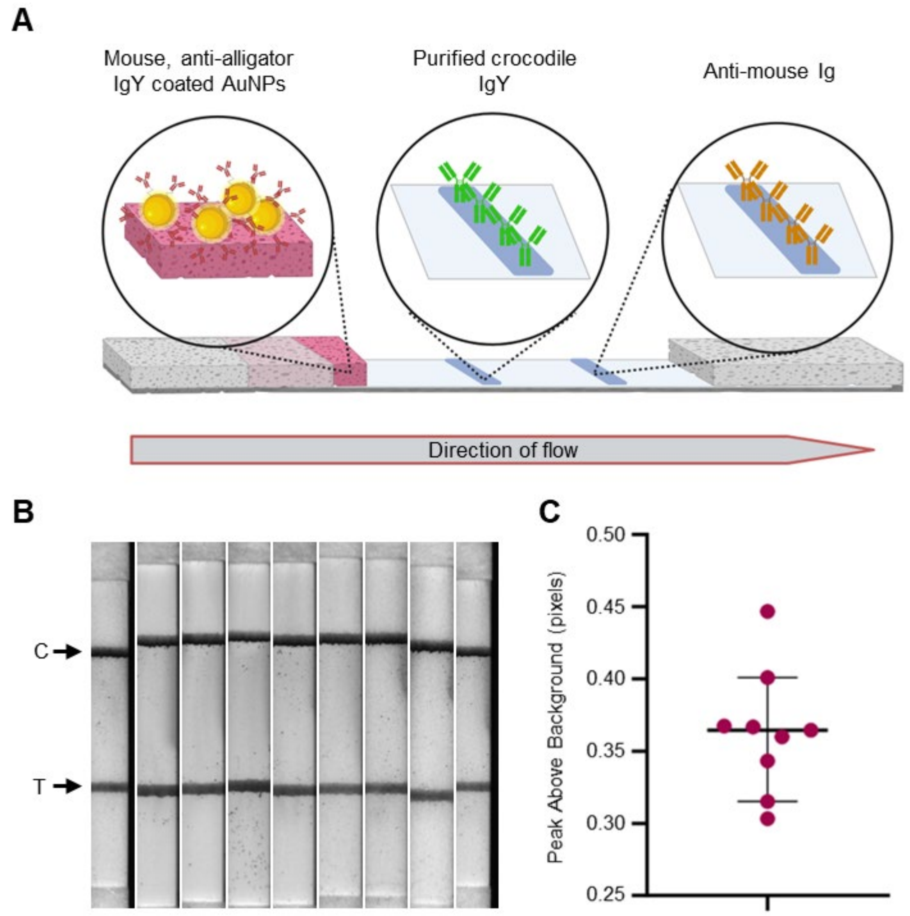


***Figure S1. Anti-alligator conjugate compatibility assessment. (A)*** *Design of LFA for assessing viability of anti-alligator IgY coated AuNPs for the detection of crocodile IgY Abs. The design features mouse, anti-alligator IgY conjugated AuNPs, purified crocodile IgY printed on the test line and an anti-mouse Ig is printed as the control line.* ***(B)*** *Visual representation of the nine strips run, indicating the binding of the anti-alligator conjugate to the crocodile Abs. T = test line and C = control line.* ***(C)*** *Nine strips were run, imaged using the Leelu reader and the results graphed. Bars indicate median and 95% confidence interval. Created in part with Biorender.com*
