## Supplementary figures and images for "Application of chimeric antigens to paper-based diagnostics for detection of West Nile virus infections of *Crocodylus porosus –* a novel animal test case"

### Supplemental Figure 1

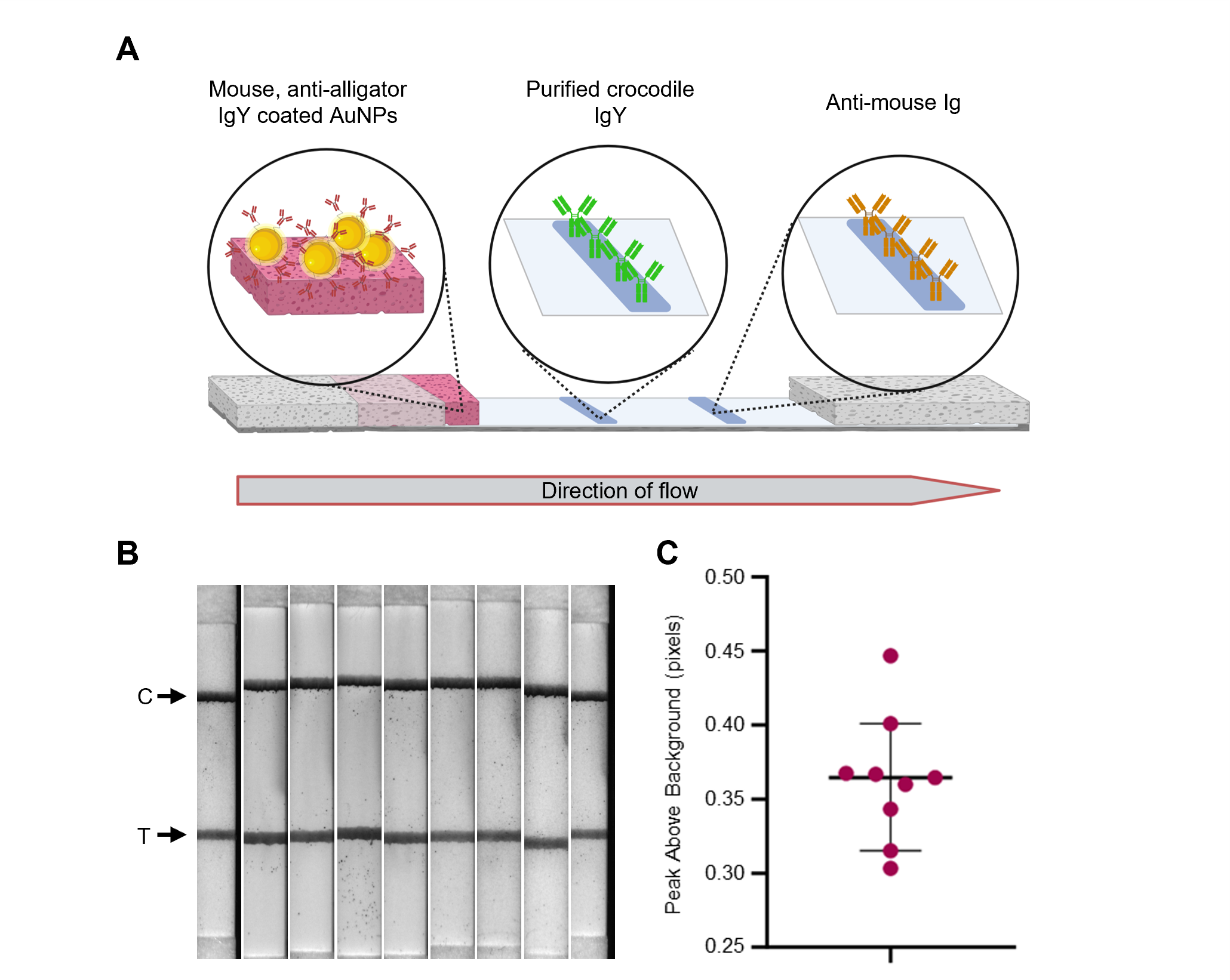

### Supplemental Table 1

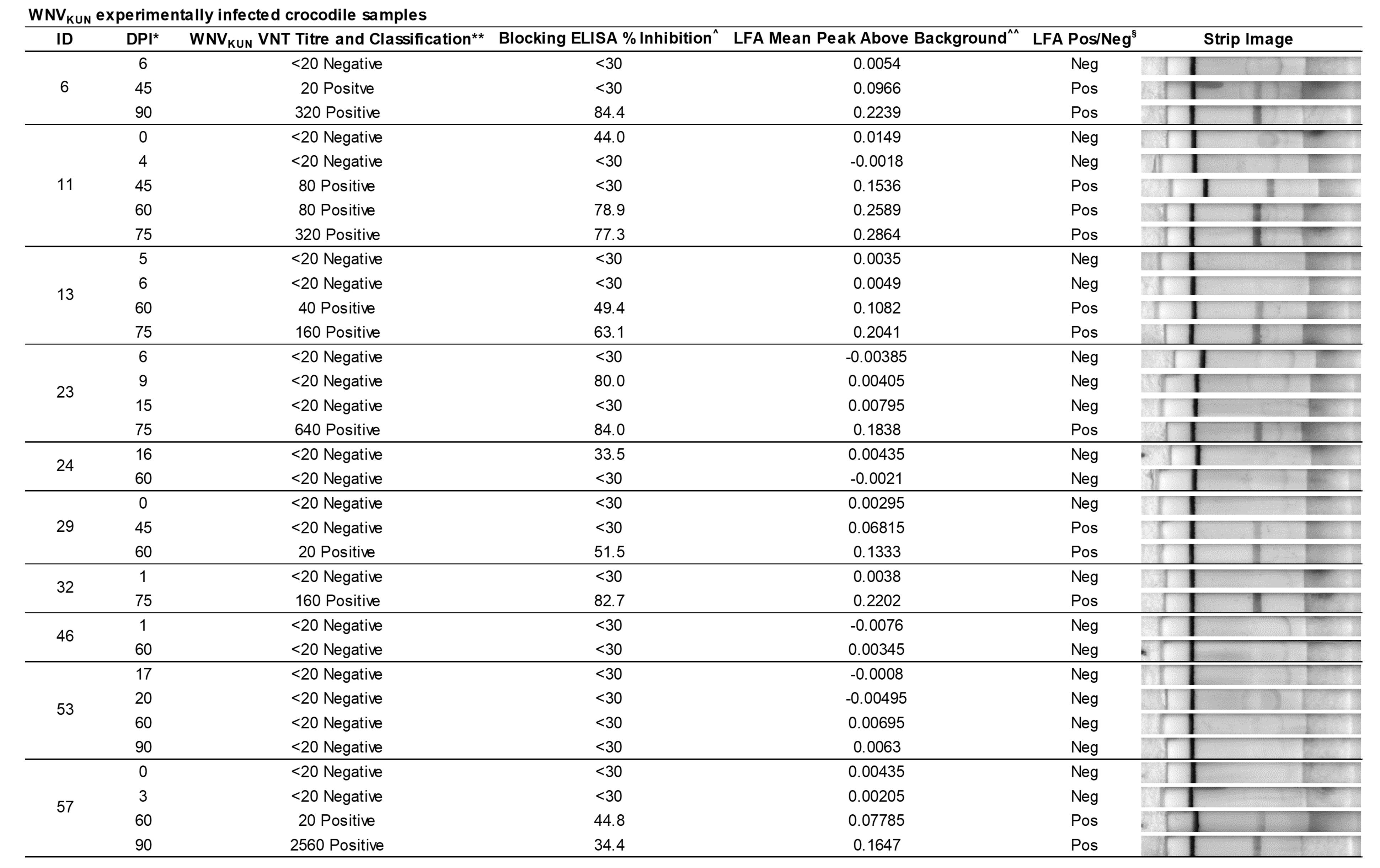

### Supplemental Table 2

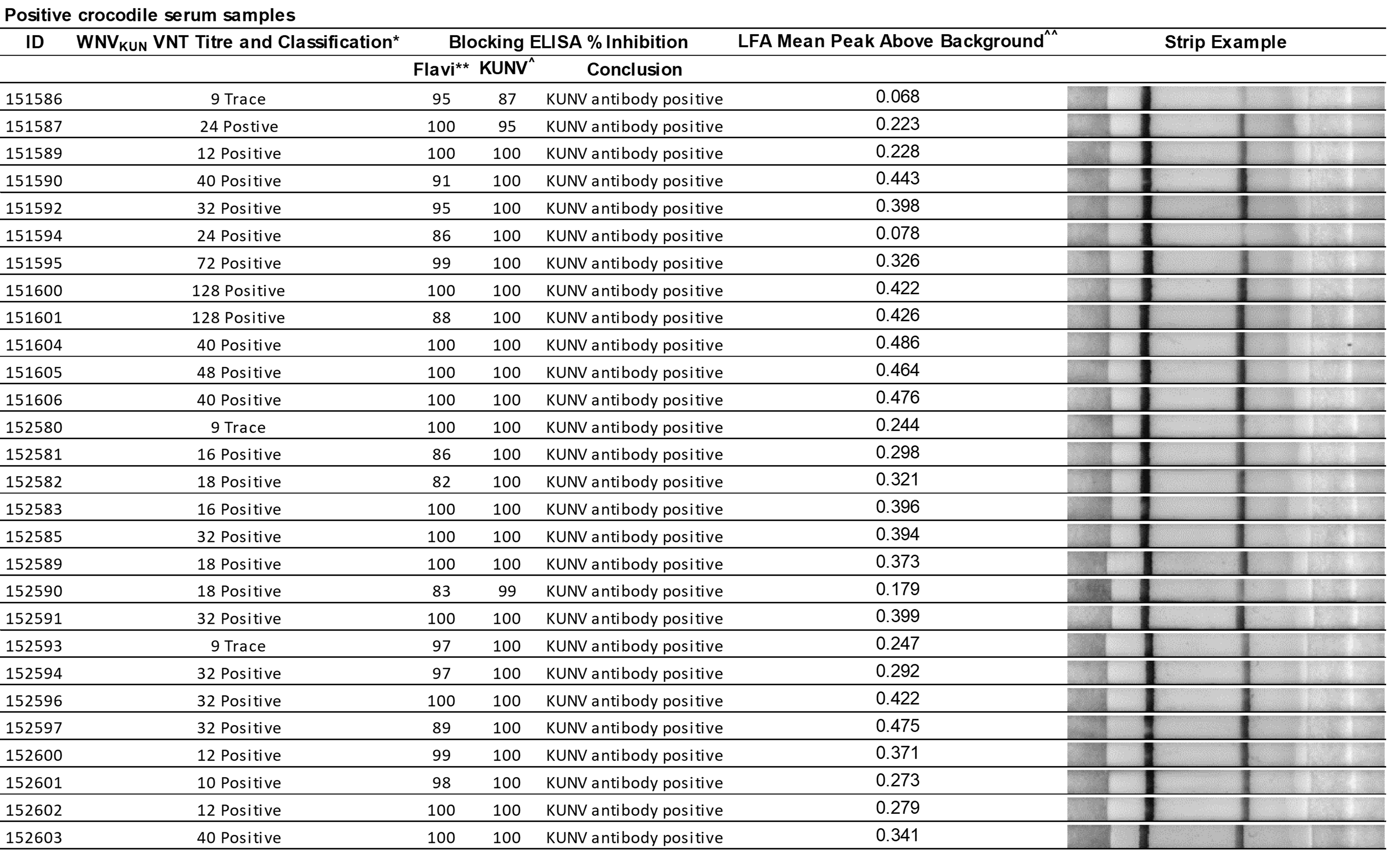

### Supplemental Table 3

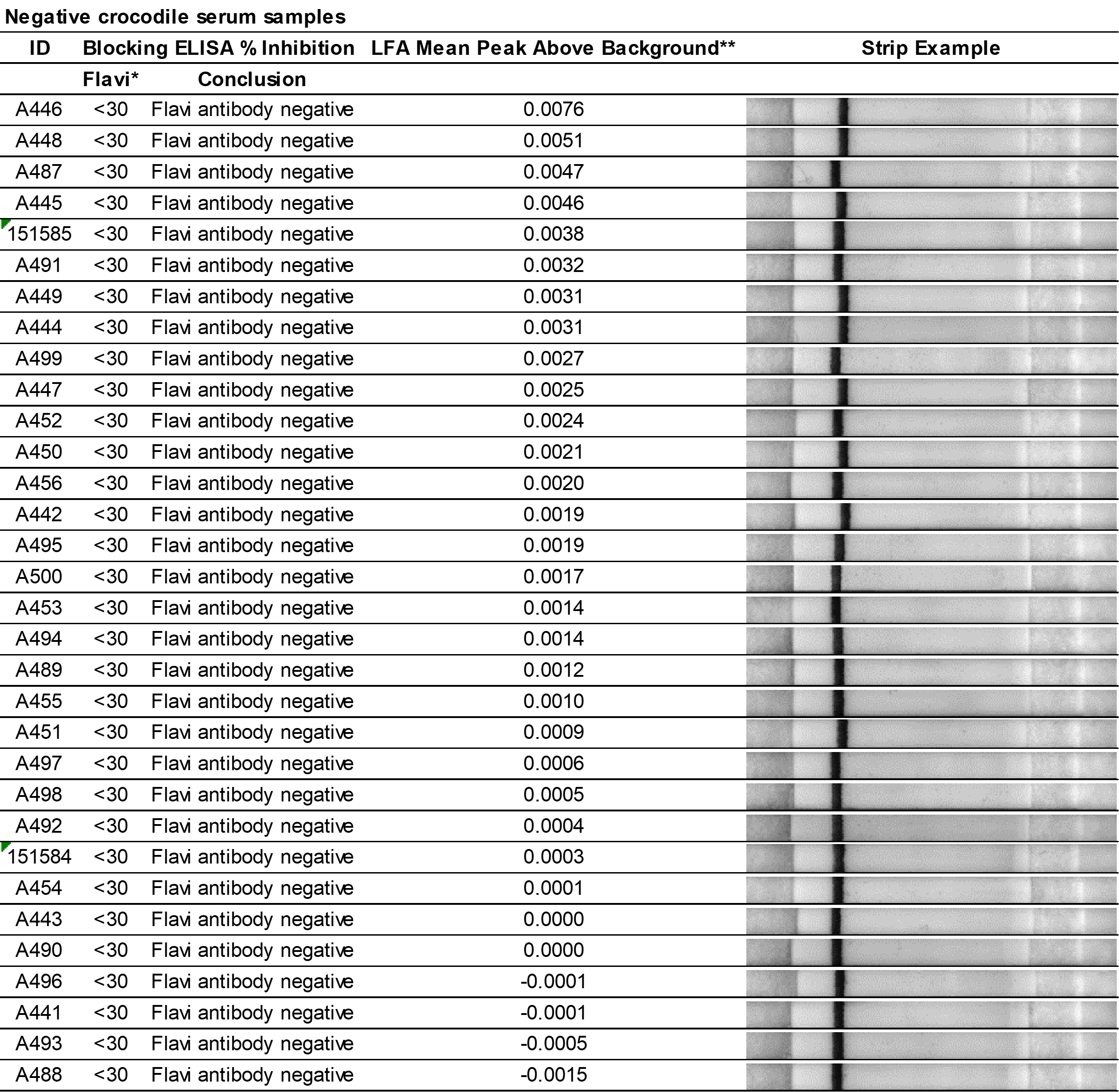
